# Absolute Winding Number Differentiates Spatial Navigation Strategies with Genetic Risk for Alzheimer’s Disease

**DOI:** 10.1101/2022.02.04.479128

**Authors:** Alexandra Badea, Didong Li, Andrei R Niculescu, Robert J Anderson, Jacques A Stout, Christina L Williams, Carol A Colton, Nobuyo Maeda, David B Dunson

## Abstract

Spatial navigation and orientation are emerging as promising markers for altered cognition in prodromal Alzheimer’s disease, and even in cognitively normal individuals at risk for Alzheimer’s disease. The different APOE gene alleles confer various degrees of risk. The APOE2 allele is considered protective, APOE3 is seen as control, while APOE4 carriage is the major known genetic risk for Alzheimer’s disease. We have used mouse models carrying the three humanized APOE alleles and tested them in a spatial memory task in the Morris water maze. We introduce a new metric, the absolute winding number, to characterize the spatial search strategy, through the shape of the swim path. We show that this metric is robust to noise, and works for small group samples. Moreover, the absolute winding number better differentiated APOE3 carriers, through their straighter swim paths relative to both APOE2 and APOE4 genotypes. Finally, this novel metric was sensitive to sex differences, supporting increased vulnerability in females. We hypothesized differences in spatial memory and navigation strategies are linked to differences in brain networks, and showed that different genotypes have different reliance on the hippocampal and caudate putamen circuits, pointing to a role for white matter connections. Moreover, differences were most pronounced in females. This departure from a hippocampal centric to a brain network approach may open avenues for identifying regions linked to increased risk for Alzheimer’s disease, before overt disease manifestation. Further exploration of novel biomarkers based on spatial navigation strategies may enlarge the windows of opportunity for interventions. The proposed framework will be significant in dissecting vulnerable circuits associated with cognitive changes in prodromal Alzheimer’s disease.

## 1 Introduction

Alzheimer’s disease (AD) directly affected 6 millions Americans in 2021, and these numbers include more than 12% of women, and 9% of men older than 65 (Alzheimer’sAssociation 2021). The disease starts before overt memory loss and difficulty thinking, but escapes detection for decades, by which time it is too late for current treatments to be effective. A strategy to overcome these limitations and to quicken the pace of discovery, is to study people at risk for AD. The largest known genetic risk factor for AD is linked to the APOE gene. Having one copy of the APOE4 allele can increase risk for late onset AD by 2 to 3 times while two copies can increase the risk by 12 times (Michaelson 2014). In contrast, the APOE2 allele is thought to decrease risk for AD, relative to control APOE3 carriers and at risk APOE4 carriers (Wu and Zhao 2016). Humanized mouse models expressing these three major human APOE isoforms (targeted replacement) (Sullivan et al. 1998) (Knouff et al. 2004) can also be used to model genetic risk for late onset Alzheimer’s disease.

Studying human populations and animal models of genetic risk for AD gives us the possibility to identify early biomarkers of AD. While the main complaints in AD are memory impairment and difficulty thinking, these are detected late in the disease process. Spatial navigation and orientation symptomatology have also been reported in AD, while the method chosen and performance in spatial strategies may provide protection against hippocampal degeneration during aging (Bohbot et al. 2007a). It has been suggested that spatial navigation impairment, in particular for allocentric and real space configurations, occurs early in the development of AD and can be used for monitoring disease progression or for evaluation of presymptomatic AD (Hort et al. 2007). Recent studies suggest that midlife *APOE4* carriers exhibit changes in navigation patterns before any detectable symptom onset (Coughlan et al. 2018).

While we know that the hippocampus plays an important role in spatial navigation, it is becoming increasingly clear that it does not act alone to determine the goal-directed navigation strategy, but in connection with circuits involving e.g. the subiculum, thalamus, cingulate cortex, fornix, hypothalamus (Bermudez-Contreras, Clark, and Wilber 2020), and the dorsal striatum. The caudate putamen circuitry is thought to convey contextual information and to help form place-reward associations (Stoianov et al. 2018), (Pennartz et al. 2011). This new information demands a shift from hippocampal centric approaches to more extended brain subnetworks. Elements of these networks may reveal differences in individuals at risk for AD, at prodromal stages, and thus provide new biomarkers.

One way to test such target circuits is through lesion studies, and those have revealed that the (dorsal) hippocampus, fornix (Eichenbaum, Stewart, and Morris 1990), striatum, basal forebrain, cerebellum and cerebral cortex lead to lower performance; and so does disconnecting regions relevant for spatial learning. Still, it is not fully understood how different anatomical network nodes are involved in the acquisition and maintenance of different types of information required for spatial navigation, and what are the relationships with the genotypes that confer risk for AD. For example, approximately 50% of young adults prefer to use a spatial strategy, while the other 50% prefer a response strategy (Iaria et al. 2003). The spatial strategy involves using relationships between landmarks, and is thought to depend on the hippocampus (Bohbot, Iaria, and Petrides 2004).The response strategy involves learning stimulus-response associations, such as a series of right and left turns from specific points in space (McDonald and White 1994), and is thought to depend on the caudate putamen. The literature supports that the dorsal striatum is involved in stimulus–response learning, while the hippocampus mediates place learning. Moreover, increased gray matter density in the caudate nucleus has been associated with less gray matter in the hippocampus and vice versa. Therefore, navigation strategies are sensitive to the predominant use of gray matter in the hippocampus and caudate (Konishi et al. 2016) memory systems. Such relationships have been shown for the gray matter (Bohbot et al. 2007b) in humans, but the role of white matter tracts in modulating performance in spatial navigation has been less explored, in particular in relation to APOE genotypes. There is a need to better understand the role of different brain networks comprised of gray matter nuclei and their white matter connections, and how they confer vulnerability to AD. The differential roles between the two memory systems relying on the hippocampus and caudate putamen, and their associated brain circuits can be characterized using fMRI or diffusion weighted MRI and tractography, and may have the potential to reveal new and early markers in APOE carriers with different genetic risk levels.

Current studies have not consistently shown hippocampal atrophy in APOE4 carriers, in the absence of overt AD pathology. Some studies reported decreased hippocampal volume in young and old cognitively normal APOE4 carriers (Crivello et al., 2010; O’Dwyer et al., 2012; Wishart et al., 2006), while other did not find hippocampal atrophy (Haller et al., 2017; Honea et al., 2009). The structural covariance of different brain regions in relation to cognitive changes in prodromal AD has been less studied, but also points to more extended networks, where structural covariance patterns indicate differences with genotype (Novellino et al. 2019). Inverse correlation between hippocampus and caudate putamen and between these regions’ gray matter and the preference for a spatial strategy has been shown (Bohbot et al. 2007b), but how these relationships are altered in relation to APOE is less known. This supports a need to investigate other regions beyond the hippocampus to understand the vulnerability of APOE4 carriers to AD (Crivello et al., 2010). It remains to be seen if extended brain circuits involved in spatial navigation may offer novel targets.

To assess spatial navigation strategies in subjects at risk is noninvasive and inexpensive. These assessments can complement more invasive molecular and mechanistic studies in animal models. The Morris Water Maze (MWM) is a popular tool to test spatial learning and memory, and navigation strategies, and was originally designed for animal tests. MWM like tests have also been designed and extended to humans, e.g. using virtual reality (Hodgetts et al. 2020). In the MWM test (Morris et al. 1982) mice are placed in a circular pool and required to swim to a hidden platform beneath the surface using cues. MWM has long been thought as a test of hippocampal function, but more recently performance has been linked to the coordinated action of regions constituting a network (Hodgetts et al. 2020). Most often the performance in the water maze is described by the escape latency, or distance swam until the animals find the hidden platform. Search strategies are less often described, and rarely quantified, e.g. as manually scored percentage of time/distance spent using different strategies such as spatial, systematic, or looping search patterns. Using such techniques has helped identify increased chaining/loopiness following parietal cortex injuries (Brody and Holtzman 2006; Tucker, Velosky, and McCabe 2018) (Brabazon et al. 2017). In this paper we introduce a novel metric to characterize the search strategy, or loopiness of the swim path, the absolute winding number, and we assess its ability to discriminate between carriers of the three major APOE alleles.

Finally, we related changes in swim path shape, or search strategy, to imaging metrics derived from high resolution, high field MRI. For our analyses we selected two regions involved in spatial navigation, the hippocampus and the caudate putamen, as well as their major connections, through fimbria and fornix on one hand, and the internal capsule on the other. We added the cerebellar white matter to examine its role in modulating the search strategy as well, although this region is frequently used as a control region in AD studies. More recently the cerebellum has emerged as also having a role in learning, and it has been suggested it may interact with the hippocampus (Babayan et al. 2017), perhaps via other regions, including the retrosplenial cortex (Rochefort, Lefort, and Rondi-Reig 2013).

Our goals were to dissect whether spatial learning and memory circuits are differentially modulated by APOE isoforms, in the absence of AD pathology, and whether female sex confers increased vulnerability. Animal behavior was assessed in the Morris water maze in mouse models that express either human APOE2, APOE3, or APOE4 alleles, to reveal the impact of APOE genotype on brain circuit vulnerability in aging/AD. In a subset of mice, we have compared how learning and memory markers relate to the hippocampus and striatum structural phenotypes, using diffusion weighted imaging to characterize morphometry through volume changes between genotypes, microstructural properties through fractional anisotropy, and connectivity through degree and clustering coefficient.

Our outcomes include factors such as behavior characteristics of spatial learning and memory, morphometry and texture based on MR imaging markers, and tractography based connectomics. We introduced a novel marker to the traditional distance measures, to characterize the complexity of the navigation strategy in the MWM, through an absolute winding number. This describes the loopiness of the swim path of mice tasked to locate a submerged platform in the Morris Water Maze, in a quantitative manner that makes it amenable to compare such strategies directly, and adds to the existing battery of MWM based metrics. We compared these behavioral and imaging markers with genotype, and sex. We build models to help distinguish how navigation strategies map to different brain regions and circuits in mice with the three major APOE alleles. Our analyses revealed that both genotype and female sex play a role, differentiating the three APOE alleles, and that the absolute winding number adds a robust and sensitive marker that may find translational applications if added to human studies evaluating genetic risk for AD.

## 2 Methods

### 2.1 Animals

To dissect how brain circuits vulnerable in aging and AD are modulated by the three major APOE isoforms, we have examined spatial learning and memory function using the Morris Water Maze test, in relation to morphometric and connectivity characteristics of the hippocampal, striatal and cerebellar circuits, as determined from diffusion weighted based MRI.

We used 12 month old humanized mice modeling genetic risk for late onset Alzheimer’s disease expressing the three major human APOE alleles (targeted replacement). Mice were homozygous for the APOE2 allele, thought to be protective against Alzheimer’s disease; APOE3, thought as the control gene variant, or APOE4, which is the major known genetic risk for late onset AD. Animals included both male and female sexes (**Table 1**).

**Table1.** Animal groups distribution by genotype, sex, and age range.

| Genotype | No Animals | Males | Female | Mean Age (months) | SD Age (months) |
| --- | --- | --- | --- | --- | --- |
| APOE2 | 13 | 8 | 5 | 12.64 | 0.70 |
| APOE3 | 17 | 6 | 11 | 12.70 | 0.98 |
| APOE4 | 24 | 13 | 11 | 12.47 | 1.43 |

### 2.2 Spatial Learning and Memory Testing

Mice were handled for 5 days prior to behavioral testing to habituate to the researchers performing the tests, and to water. Spatial learning and memory were assessed using the Morris Water maze paradigm, similar to (Badea et al. 2019). The MWM tests a mouse’s spatial memory and learning based on their preference for standing on solid ground, as opposed to swimming. Mice were trained for 5 days in a circular swimming pool, filled with water rendered opaque using nontoxic white paint. The pool has 150 cm diameter, and behavior in the pool was tracked with a ceiling-mounted video camera, and the ANY-maze (Stoelting, Wood Dale, IL, United States) software. Four trials were administered each day, in blocks of 2, separated by 30 minutes, and trials ended after 1 min maximum. Each trial consisted of placing the mouse into the water at one of four different starting positions, one in each quadrant. The quadrant order was varied each day. Mice could use visual cues to orient themselves, and to find refuge on a platform submerged ~1.5 cm underneath the water. Because of their aversion to swimming and the consistent placement of the platform, mice are expected to learn that the platform is located in the same position relative to directional cues and locate it more quickly over time. We assessed learning by measuring the distance mice needed to swim to reach the platform, and the distance it swam in the pool, as well as the percent swim distance in the target quadrant in which the platform is located. If mice were unable to locate the platform within the allotted time of 1 minute, they were guided to the platform and allowed to remain there for 10 s. Probe trials were conducted on days 5, 1 h after the last training trial, and then on day 8. During the probe trials the submerged platform was removed and mice were given 1 min to swim in the pool. Navigation strategies and efficiency were assessed using traditional measures such as the total swim distance, and the distance spent in each of the quadrants.

### 2.3 Absolute Winding Number

In addition to the distance metrics traditionally used to describe behavior in the Morris Water Maze paradigm, we characterized the swim path using a novel metric, the absolute winding number. This is derived from the well-known winding number in mathematics, is positive-valued and characterizes the shape of the swimming trajectory, as defined below.

#### 2.3.1 Winding Number

Consider a continuous curve 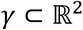 defined by the equation

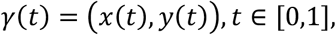

where *x* = *x*(*t*), *y* = *y*(*y*) are continuous functions, and *γ* is a closed curve if *γ*(0) = *γ*(1).We assume *γ* does not pass through the origin (0,0), and reparametrize the curve in polar coordinates as:

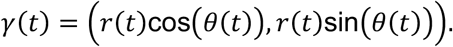

The winding number of *γ* is then defined as

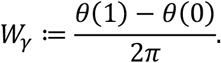

For any continuous closed curve, its winding number is always an integer, and measures the total number of times that curve travels counterclockwise around the origin. The winding number is an important object of study in differential geometry, complex analysis and algebraic topology.

#### 2.3.2 Absolute Winding Number

Our motivation in considering the winding number is to obtain a summary of how much each animal’s movement trajectory deviates from a direct path. However, the winding number is not directly useful as such a summary for three reasons: (1) the animal tracking data do not directly provide *γ*(*t*), instead yielding points along the curve at a finite number of times; (2) the curves are not closed as the animals do not return to their starting locations; (3) the movement is not expected to be consistently counterclockwise and may change between clockwise and counterclockwise. To address these limitations and obtain a more appropriate measure, we propose an Absolute Winding Number (AWN):

#### Definition 1.

*Let* 0 ≤ *t*_0_ < *t*_1_ < … < *t_n_ and γ_i_* = *γ*(*t_i_*) = (*x*(*t_i_*), *y*(*t_i_*)) *i*, = 0, …, *n be discrete points on a curve γ, with n*≥3. *Assume for any* 0 ≤ *i* ≤ *n* – 1, *γ_i_* ≠ *γ*_*i*+1_, *the Absolute Winding Number (AWN) of γ, denoted by* 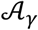, *is defined as* 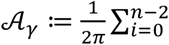 arccos 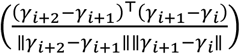

The assumption *γ_i_* ≠ *γ*_*i*+1_ means that the animal does not remain at exactly the same location between measurement times. The proposed 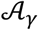 is always non-negative, is not necessarily an integer, and provides a measure of the degree of deviation of the movement trajectory from a straight line.

##### Proposition 1.

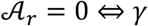 *is a straight line*.

#### 2.3.3 Continuous Absolute Winding Number

The AWN in Definition 1 depends on the sampling times *t_i_*, but provides an estimate approximating a continuous AWN (CAWN), which we define below. Let 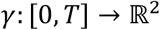 be a plane curve, and consider the unit tangent field along *γ*, denoted by *X*: [0, *T*] → *S*^1^, where *S*^1^ is the unit circle, as

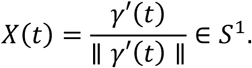

Then we represent *X* by the circular angle curve *θ*, that is:

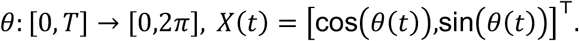

The continuous AWN is the length of the curve *θ*:

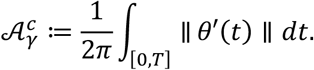

We formulate the continuous analogue of Proposition 1.

##### Proposition 2.

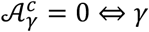 *is a straight line*.

*Proof*. 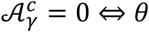 is a constant curve on *S*^1^ ⇔ *X* is a constant vector field ⇔ *γ* is a straight line.

The following Proposition implies that AWN is a discretization of CAWN: as the sample times *t_i_* get closer and closer together, AWN converges to CAWN:

#### Proposition 3.

Letting Δ*t* = sup_*i*_ |*t*_*i*+1_ – *t_i_*| to be the maximum difference between times, then 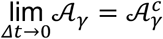.

*Proof*. Given a partition 0 ≤ *t*_0_ < *t*_1_ < … *t_n_* = *T*, observe that

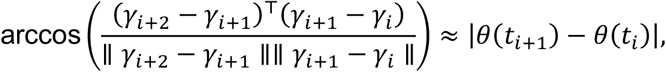

then

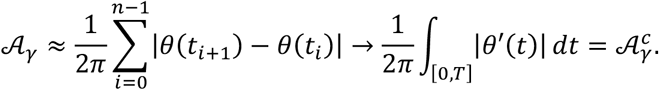

#### 2.3.4 Robustness of the Absolute Winding Number

To characterize errors in tracking movement, suppose we observe *ξ_i_* = *γ_i_* + *ϵ_i_* with noise *ϵ_i_*~*N*(0, *σ*^2^*Id*), where *Id* is the two-dimensional identity matrix. We show in Theorem 1 that the estimate of AWN based on noisy data, 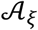, is close to the true 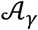 with high probability. This demonstrates robustness of the AWN.

##### Theorem 1.

*Assume there exists l*_0_ > 0 *and* 0 < *φ*_0_ < 1 *such that* ∥*γ*_*i*+1_ – *γ_i_*∥ ≥ *l*_0_ *for* 0 ≤ *i* ≤ *n* – 1 *and* 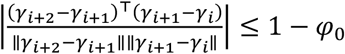 *for* 0 ≤ *i* ≤ *n* – 2, *then for any* 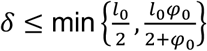, *with probability* 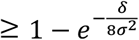, we have the following bound:

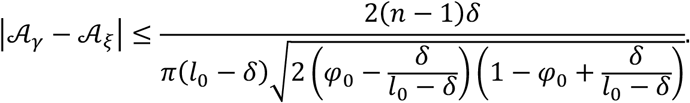

*Proof*. By the definition of AWN and triangular inequality, it suffices to consider a single time interval, that is, to compare arccos 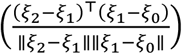 with arccos 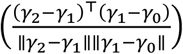. To simplify the notation, let *u*_2_ = *γ*_2_ – *γ*_1_ and *u*_1_ = *γ*_1_ – *γ*_0_, *η*_2_ = *ϵ*_2_ – *ϵ*_1_, *η*_1_ = *ϵ*_1_ – *ϵ*_0_. Then we want to analyze:

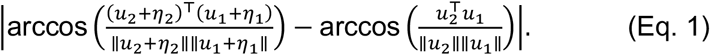

By the triangular inequality again, (Eq. 1) is upper bounded by *A + B* where

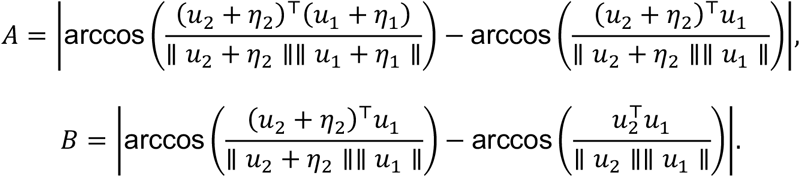

By symmetry, it suffices to bound either term so we focus on B. Let *f*(*η*) ≔ arccos(*ξ_η_*) ≔ arccos 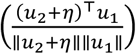, then *B* = |*f*(*η_2_*) – *f*(0)|, where *η*_2_ ~ *N*(0,2*σ*^2^*Id*). Since *η*_2_ ~ *N*(0,2*σ*^2^*Id*), 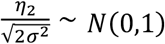 and 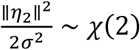 then by the tail probability of *χ*(2), for any 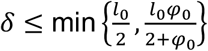, with probability at least 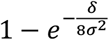, ∥*η*_2_∥ ≤ *δ*. Then with high probability, we have the following:

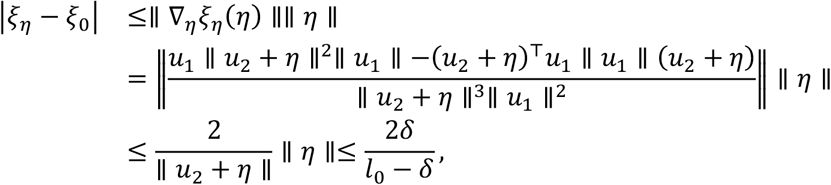

where the last inequality follows from the assumption ∥*u*_2_∥ ≥ *l*_0_ and 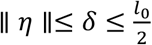. As a result,

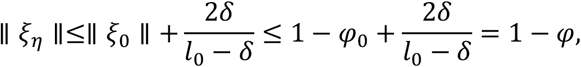

where 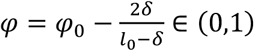 since 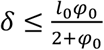. Then we observe that the gradient of *f* with respect to *η* is

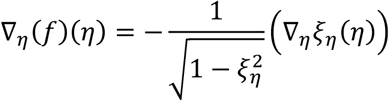

Hence

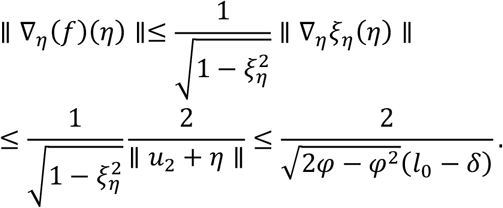

Finally, we can show:

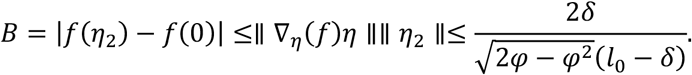

Combining the above inequalities, we have:

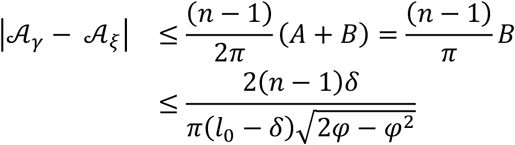

with probability at least 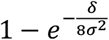 for any 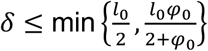.

The above Theorem implies that the larger *l*_0_ and *φ*_0_, the more robust the AWN. As a result, in practice, if two consecutive observations *γ_i_* and *γ*_*i*+1_ are too close or the inner product between the normalized 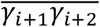 and 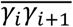 is very close to 1, then we can remove *γ_i+_1* to reduce the impact of random noise. This is not surprising since if two consecutive observations are almost identical, then tiny noise will result in huge errors in the angle. Similarly, if the two vectors are almost co-linear, the noise will contribute more to the true angle, which comes from the fact that the arccos function has infinite derivative at ±1.

### 2.4 Imaging and Associated Metrics

Diffusion weighted imaging was done using a 9.4T high field MRI, with a 3D SE sequence with TR/TE: 100 ms /14.2 ms; matrix: 420 x 256 x 256; FOV: 18.9 mm x 11.5 mm x 11.5 mm, 45 μm isotropic resolution, BW 62.5 kHz; using 46 diffusion directions, 2 diffusion shells (23 at 2000, and 23 at 4000 s/mm^2^); 5 non diffusion weighted (b0). The max diffusion pulse amplitude was 130.57 Gauss/cm; duration 4 ms; separation 6 ms, 8-fold compressed-sensing acceleration (Wang et al. 2018) (Robert Anderson 2018) (Uecker M; Ong F; Tamir JI; Bahri 2015). Diffusion data were reconstructed using DIPY (Garyfallidis et al. 2014) with Q-ball Constant Solid Angle Reconstruction, producing ~2 million tracts. We have used pipelines implemented in a high-performance computing environment, to segment the brain in sub regions (Anderson et al. 2019). We focused on a subset including the hippocampus, caudate-putamen, and their main connections, the fimbria and fornix, and the internal capsule, as well as the cerebellar white matter. For these regions we calculated features including volume and microstructural properties like fractional anisotropy (FA), to reconstruct tracts and build connectivity matrices. We used the Brain Connectivity toolbox (Rubinov and Sporns 2010) to calculate degree of connectivity (DEG) and clustering coefficient (CLUS) for the hippocampus and caudate putamen and associated fiber tracts, including fimbria (fi) and fornix (fx) for the hippocampus (Hc), and internal capsule (ic) for the caudate putamen (CPu), respectively as well as the cerebellar white matter (cbw).

### 2.5 Statistical Analyses

Statistical analyses were conducted in R to build linear models, and apply ANOVA analyses to determine the effects of genotype and sex on the behavioral markers of interest, including the total swim distance, normalized swim distance in the target quadrant, and the absolute winding number introduced above. The ANOVA analyses were followed by post hoc tests (using Sidak adjustments), and p <0.05 was considered significant. We similarly analyzed the regional volumes and FA, as well as the degree of connectivity and clustering coefficient. We used the emtrends function (emmeans R package), and evaluated linear models to relate behavioral metrics to the imaging and connectivity markers to understand if they influence the AWN, and if different genotypes/sexes use preferentially different circuits.

## 3 Results

### 3.1 Swim Paths

A qualitative analysis revealed that swim paths for selected individuals from each of the three genotypes, differed in length and shape for the learning trials and the probe tests administered in day 5, 1 hour after the trials ended, and on day 8. The last trial of day 1 is shown in **Figure 1**, since animals are likely to swim for ~1 minute during the first day (A), and this is the same duration as in the probe tests, shown in (B) and (C).

**Figure 1.**
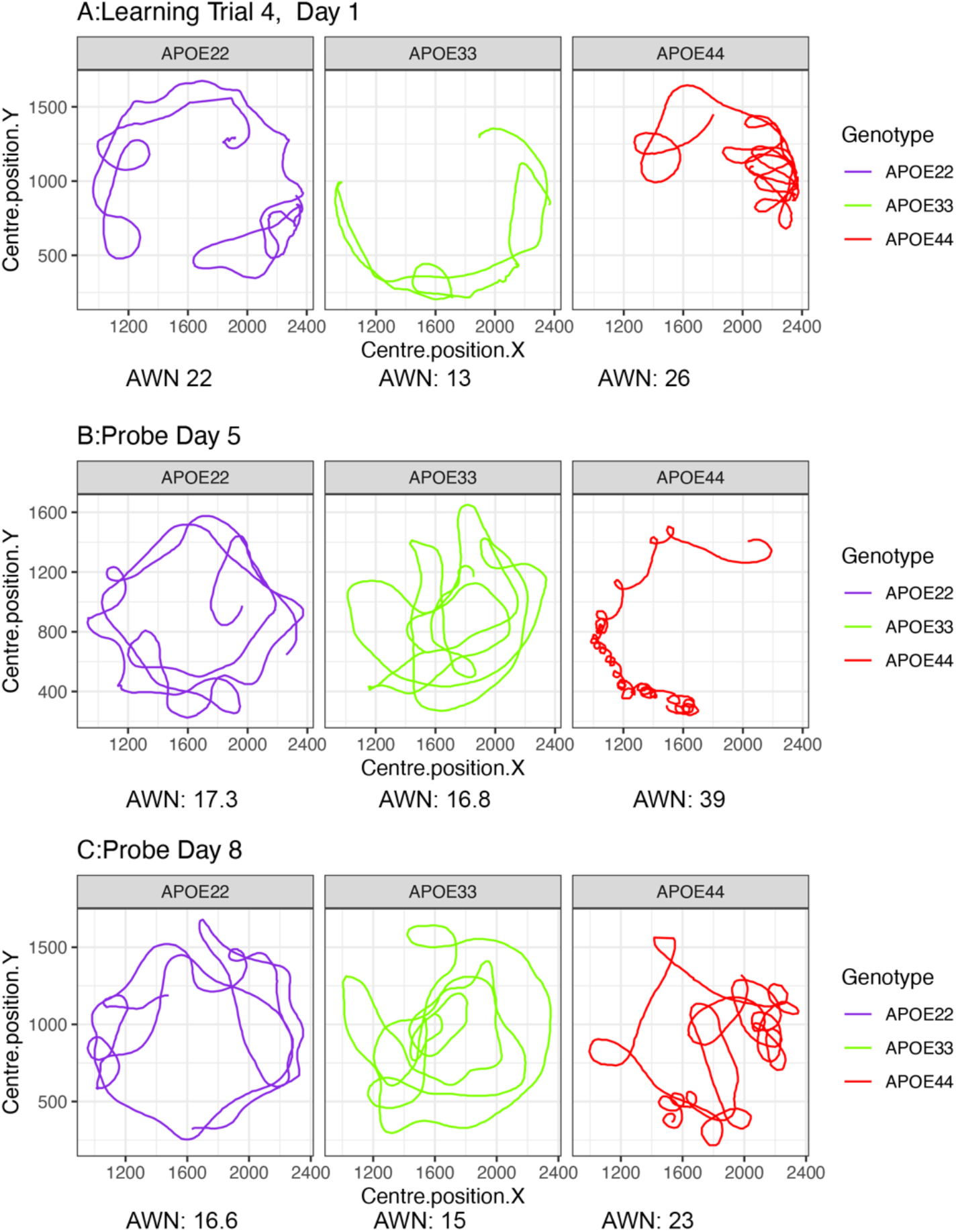
Examples of swim paths shapes for animals with APOE2, APOE3, and APOE4 genotypes. Qualitative observations suggest that swim paths differed not just in length, but also in shape. Trajectories are presented for the last trial in day 1 (A), probe in day 5 (B) and probe in day 8 (C). We chose animals illustrating medium (APOE2: learning=22, d5=17.3; d8=16.6), medium-small (APOE3: learning=13, d5=16.8, d8=15), and large winding numbers (APOE4: learning=26, d5=39; d8=23). APOE22: homozygous for APOE2; APOE33: homozygous for APOE3; APOE44: homozygous for APOE4.

### 3.2 Learning Trials

An ANOVA analysis for the total distance to the platform (**Figure 2 A, B**) revealed a significant effect of time (F(4,240)=128.2, p=2.2*10^-16^), genotype (F(2,262)=15.9, p<3.2*10^-7^), and a significant interaction of genotype by sex (F(2,262)=4.2, p=0.02). Post hoc tests indicated that differences within female groups were significant for APOE2 versus APOE3 genotypes (t=4.1, p=1.9*10^-4^), as well as between APOE3 and APOE4 genotypes (t=-4.9, p<10^-5^). The differences within male groups were significant for APOE2 versus APOE3 genotypes (t=2.5, p=0.03), as well as between APOE2 versus APOE4 genotypes (t=2.5, p=0.04). Differences were larger between females, and only APOE4 females were different relative to APOE3 females, but this was not true for males of the same genotypes. Sex differences were significant for APOE3 mice (t=-2.2, p=0.03), and showed a trend for APOE4 mice (t=1.75, t=0.08).

**Figure 2.**
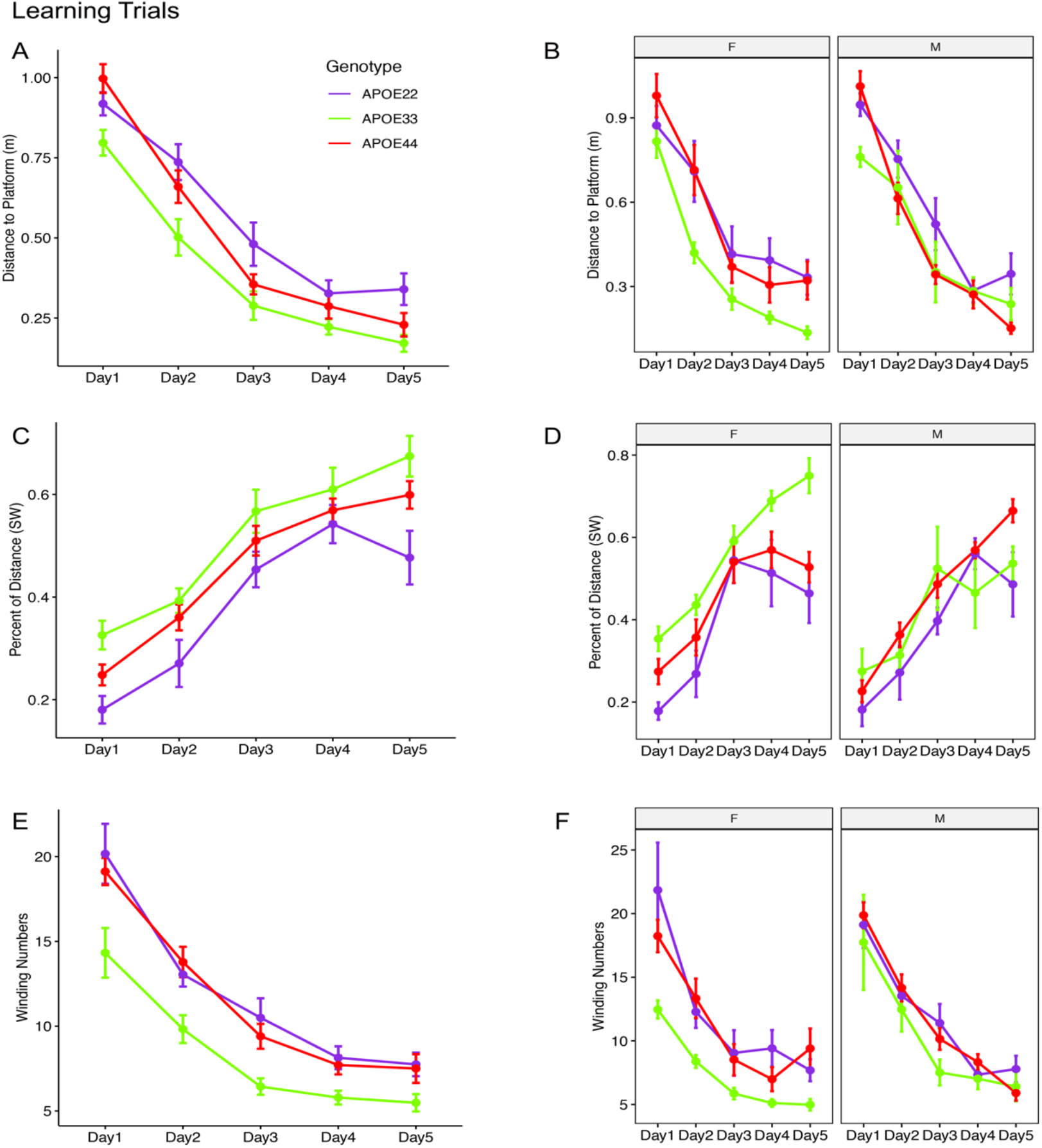
Learning trials. Mice swam shorter distances over the 5 testing days until reaching the hidden platform, indicating that they were learning. Meanwhile, the percentage time swam in the target quadrant increased with time. The absolute winding number clearly discriminated the APOE3 mice relative to APOE2 and APOE4 carriers, which used more similarly shaped trajectories. The effects were larger in females across the 5 days. F: female; M: male. Graphs show mean±standard error.

An ANOVA analysis for the normalized distance swam in the target (SW) quadrant (**Figure 2 C, D**) revealed that there was a significant effect of time (F(4,232)=69.8, p=2.6*10^-16^) and genotype (F(2,232)=13.6, p<2.6*10^-6^), a significant interaction of genotype by sex (F(2,232)=8.5, p=0.0003), and a trend for the interaction of genotype by sex by time (F(8,232)=1.9, p=0.06). Post hoc tests indicated that differences within female groups were significant for APOE2 versus APOE3 genotypes (t=-4.9, p<10^-4^), as well as between APOE3 and APOE4 genotypes (t=4.5, p<10^-4^). Post hoc tests indicated that differences within male groups were significant for APOE2 versus APOE4 genotypes (t=-2.5 p=0.03). Sex differences were significant for APOE3 mice (t=4.8, p=2.2*10^-6^). Differences between APOE3 and APOE4 mice could thus be largely attributed to differences in females.

A qualitative evaluation of the absolute winding number indicated more similar swim trajectories between APOE2 and APOE4 mice, and a clear demarcation relative to APOE3 mice. Moreover, these differences appeared clearer in females. An ANOVA analysis (**Figure 2 E, F**) revealed a significant effect of time (F(4,240)=86.9, p=2.2*10^-16^), and genotype (F(2,240)=24.15, p<2.8*10^-10^), as well as a significant effect of sex, that was not captured by the previous metrics (F(1,240)=4.8, p=0.03). There was also a significant interaction of genotype by sex (F(2,240)=3.9, p=0.02), while the time by sex interaction was characterized by F(4,240)=1.7, p=0.16. Post hoc tests indicated that differences within females were significant for APOE2 versus APOE3 genotypes (t=5.4, p<10^-5^), as well as between APOE3 and APOE4 genotypes (t=-5.7, p<10^-5^). Differences within males were not significant. Thus, differences in the shape of the trajectories were explained by females. Sex differences were significant for APOE3 mice (t=-3.5, p=0.0006).

The absolute winding number better discriminated APOE3 mice relative to APOE2 and APOE4 mice, which performed more similarly in terms of their spatial navigation strategy.

### 3.3 Probe Trials – Long Term Memory

An ANOVA analysis for the total distance swam during the probe trial (1 minute) administered on day 5 (**Figure 3**), one hour after the last learning trial did not detect significant differences.

**Figure 3.**
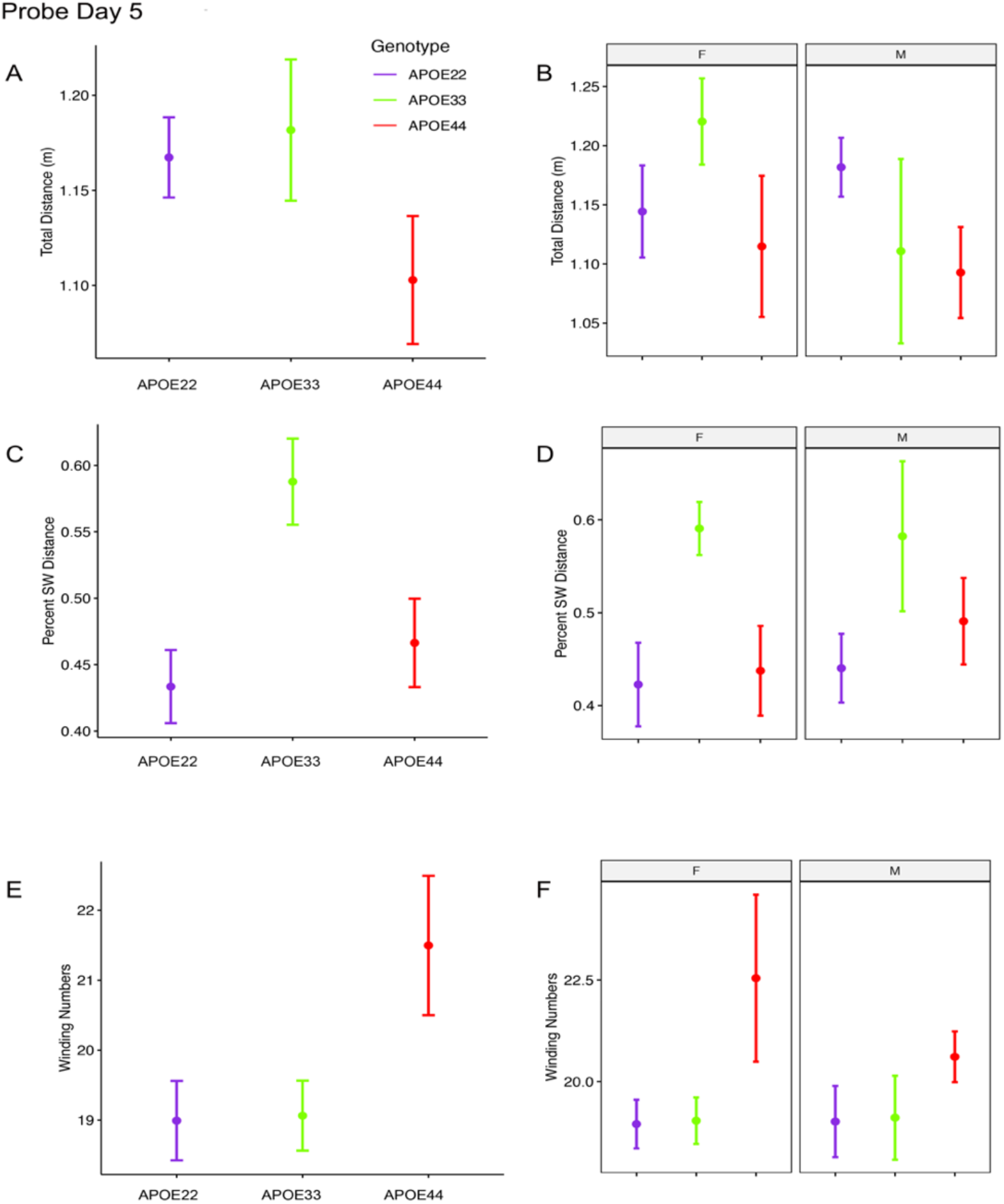
Probe trials 1 hour after ending the learning trials. Long term memory tested one hour after the end of learning trials indicated that APOE4 mice swam less than APOE2 and APOE3 mice, and the data suggested a “dose dependent” genotype effect in males. APOE3 mice spent most of their swimming in the target quadrant (~80%), while APOE2 and APOE4 mice spent (~50%) of their swimming in the target quadrant, but the differences between males and females were not significant. APOE2 and APOE4 mice were more similar, while significant differences were noticed between APOE2 and APOE3 mice, as well as between APOE3 and APOE4 female mice. The shape of the swim path, described by the absolute winding number showed similarities between APOE2 and APOE3 mice, but higher loopiness for APOE4 mice. These differences were largest for females. F: female; M: male. Graphs show mean±standard error.

The percent distance swam in the target quadrant during this probe on day 5 showed a significant effect of genotype (F(2,48)=4.5, p=0.02). Post hoc tests within groups of females (Sidak corrected) indicated significant differences within females showed a significant difference between APOE3 and APOE4 mice (t=2.5, p=0.04). Differences within groups of male mice were not significant.

The absolute winding number during day 5, testing long term memory, identified borderline significant differences due to APOE genotype (F(2,48)=3.02, p=0.06). These differences were found between females of APOE4 genotypes relative to APOE3 genotypes (t=-2.3, p=0.07). Differences within groups of male mice were not significant.

An ANOVA analysis for the total distance swam during the 1 minute of the probe trial administered on day 8 (**Figure 4**), 3 days after the last learning trial, indicated a significant effect of sex (F(1,47)=7.3, p=0.01). Post hoc tests within groups of males (Sidak corrected) indicated borderline differences between APOE2 and APOE4 genotypes (t=2.36, p=0.06). Differences between males and females of the same genotypes were found for APOE4 mice (t=2.8, p=0.007).

**Figure 4.**
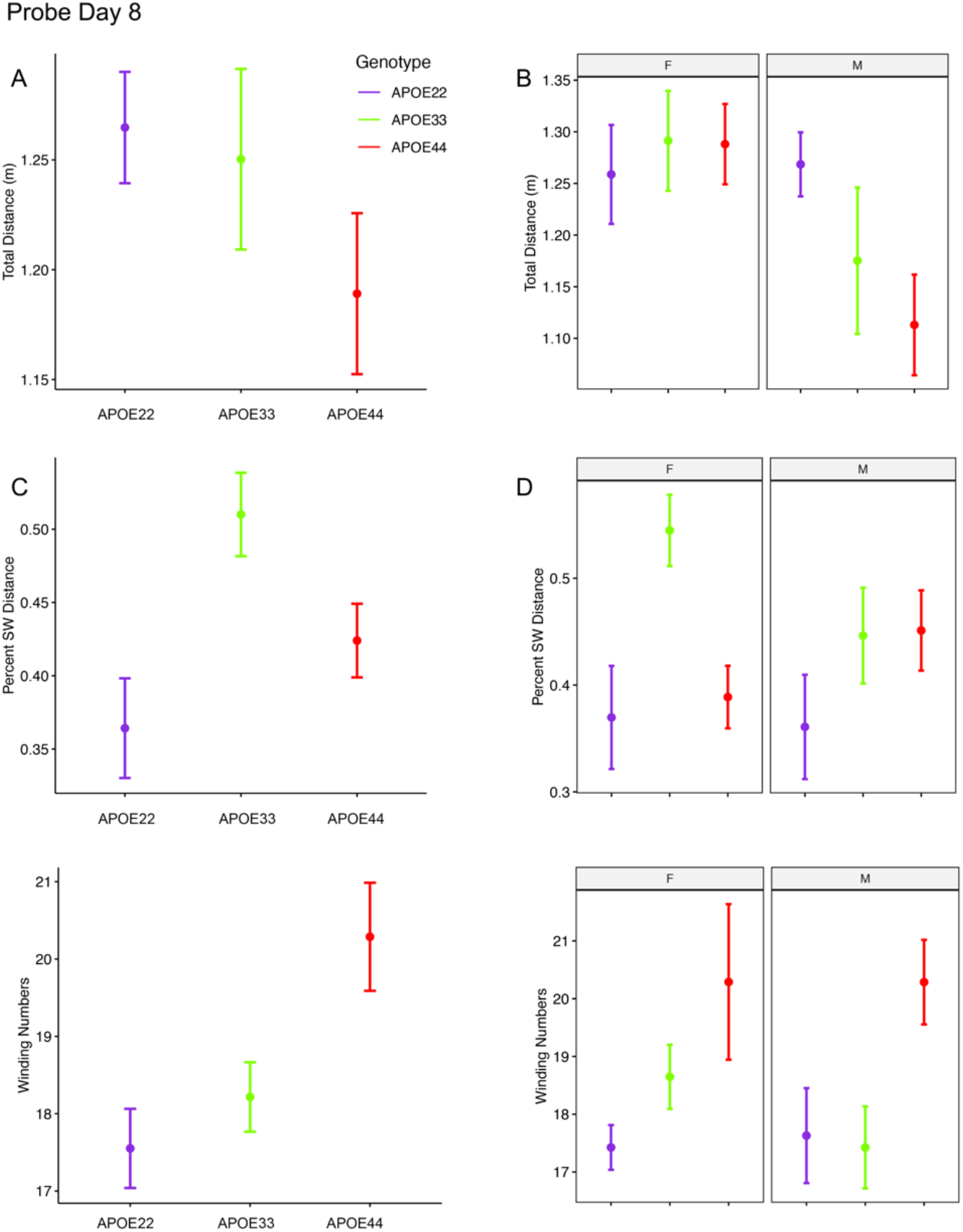
Probe trials 3 days after ending the learning trials (mean±standard error). A. The largest effects in terms of total distance were seen in males, where APOE4 mice swam the shortest distances. Our analysis does not capture stops when animals may orient themselves. **B.** The percentage time swam in the target quadrant was largest in APOE3 mice relative to APOE2 and APOE4 mice. This effect was driven largely by females, while male mice with APOE2 genotype spent less time in the target quadrant relative to other mice, and APOE3 and APOE4 mice performed similarly. A dose dependent effect was apparent in the absolute winding number for all genotypes, and this was reflected mostly in females. Male mice with APOE2 and APOE3 genotypes used similar strategies, females with APOE2 genotypes having smaller winding numbers. APOE4 males had loopier swim trajectories relative to both APOE2 and APOE3 mice, which had similar trajectories. F: female; M: male. Graphs show mean±standard error.

The percent distance swam in the target quadrant during this probe on day 8 showed a significant effect of genotype (F(2,47)=5.0, p=0.01), and a trend for the interaction of genotype by sex (F(2.47)=2.3, p=0.1). Post hoc tests within groups of females (Sidak corrected) indicated significant differences between APOE2 and APOE3 mice (t=-2.7, p=0.03), and between APOE3 and APOE4 (t=3.1, p=0.01). Differences within groups of male mice were not significant. Differences between males and females of the same genotypes showed a trend only, for APOE3 mice (t=1.7, p=0.1).

The winding number for the probe in day 8 showed a significant effect of genotype (F(2,47)=5.3, p=0.008). While differences within females were not significant, our data suggests a “dose” effect APOE2<APOE3<APOE4. Post hoc tests within groups of males showed borderline differences between APOE2 and APOE4 mice (t=-2.2, p=0.08) mice, and between APOE3 and APOE4 mice (t=-2.2, p<0.09).

Thus, the absolute winding numbers indicated more complex trajectories for APOE4 mice relative to APOE3 and APOE2 mice.

### 3.4 MRI Correlates of Spatial Navigation

As both the hippocampus and caudate putamen have been involved in spatial navigation, we examined imaging markers corresponding to changes in navigation strategies based on volume, fractional anisotropy, and structural connectivity (**Figure 5**) of these major gray matter regions, and their main white matter connections, i.e. fimbria, and fornix for the hippocampus, and the internal capsule for the caudate putamen. We have also examined the cerebellar white matter due to its less understood role, its involvement in spatial navigation, and potential hippocampal cerebellar connections (Rochefort, Lefort, and Rondi-Reig 2013).

**Figure 5.**
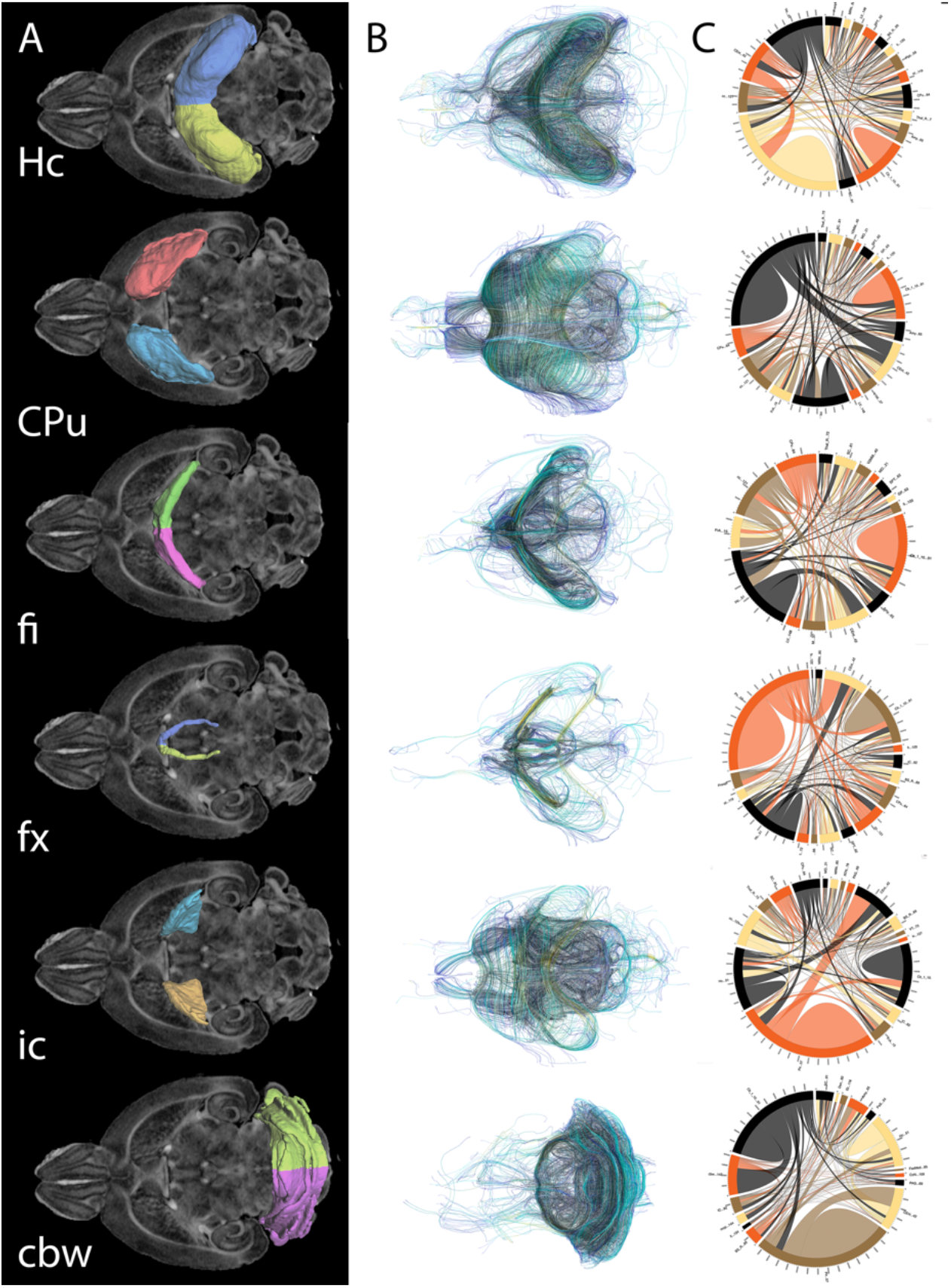
Regions of interest for spatial navigation and their MRI associated metrics. We segmented selected brain regions involved in spatial navigation, including the hippocampus (Hc), caudate putament (CPu), and their major connections (fimbria: fi, and fornix: fx; and internal capsule: ic, respectively), to which we added the cerebellar white matter, and we have measured their volumes (A). These regions were characterized by diffusion based measurements, which characterize microstructure through texture, and may vary along tracts (such as fractional anisotropy) (B). Finally we characterized their connectivity with other brain regions (C). Abbreviations and region indices correspond to the CHASS atlas (Calabrese et al. 2015),(Anderson et al. 2019) and Paxinos mouse brain atlas: Hc: hippocampus; CPu: caudate putament, fi: fimbria, fx: fornix, ic: internal capsule, cbw: cerebellar white matter.

MRI regional metrics for all three genotypes are shown in **Figure 6**, and summarized in **Table 2**.

**Figure 6.**
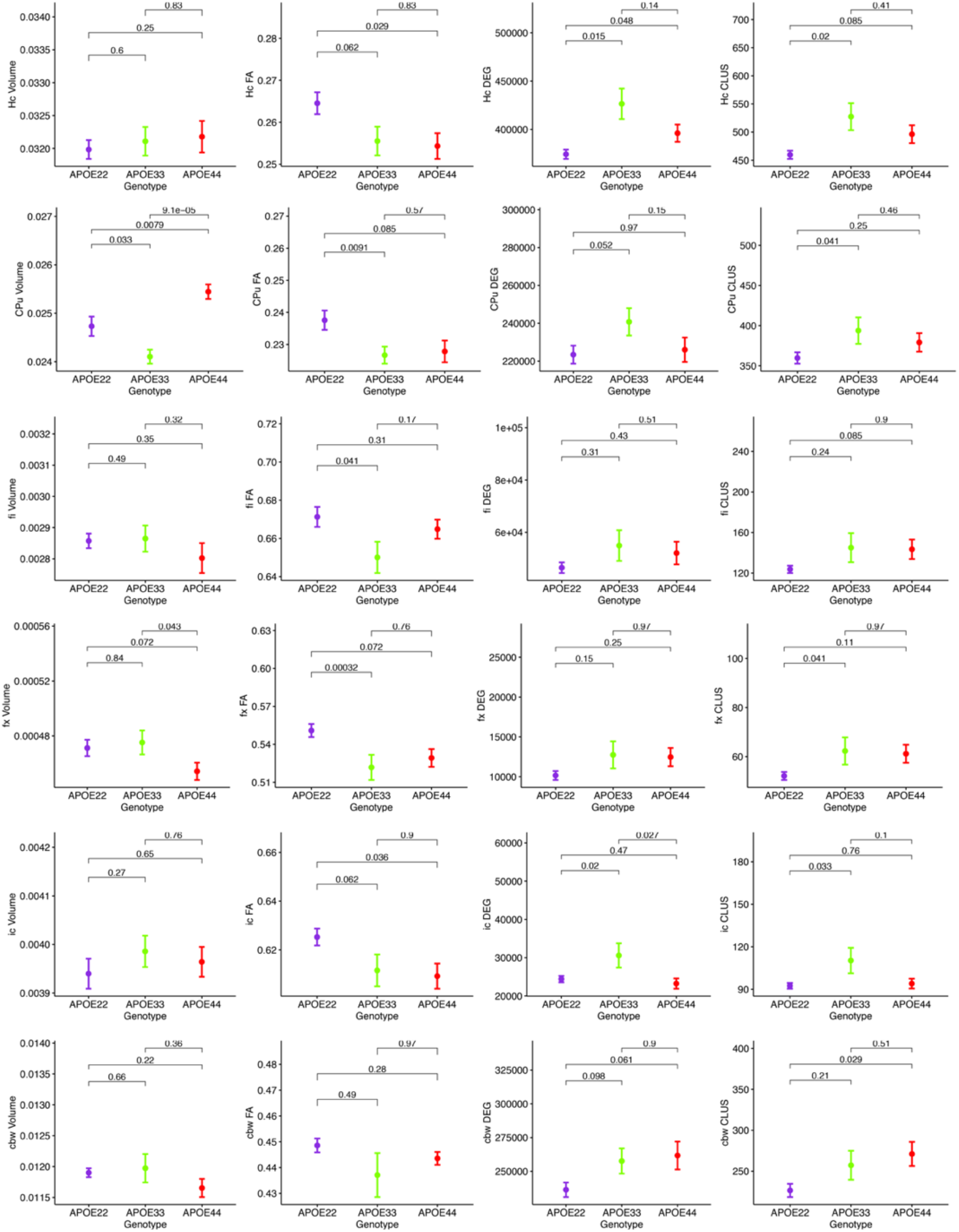
Imaging and network markers for volume, FA, degree of connectivity and clustering coefficient showed APOE genotype differences. Graphs show mean±standard error.

**Table 2.** APOE genotype differences in MRI metrics i.e. volume, fractional anisotropy (FA), degree of connectivity (DEG), and clustering coefficient (CLUS) for regions of interest including: hippocampus (Hc), caudate putament (CPu), fimbria (fi), fornix (fx), internal capsuel (ic), and cerebellar white matter (cbw).

| Region | Volume |  | FA |  | DEG |  | CLUS |  |
| --- | --- | --- | --- | --- | --- | --- | --- | --- |
|  | F | p | F | p | F | p | F | p |
| Hc | 0.43 | 0.63 | 4.11 | 0.03* | 6.9 | 0.005* | 4.34 | 0.03* |
| CPu | 13.82 | 0.0001* | 3.97 | 0.03* | 2.03 | 0.15 | 2.04 | 0.15 |
| fi | 0.86 | 0.43 | 2.92 | 0.07 | 1.12 | 0.53 | 1.62 | 0.22 |
| fx | 3.03 | 0.07 | 5.27 | 0.01* | 1.56 | 0.23 | 2.3 | 0.12 |
| ic | 0.46 | 0.62 | 5.51 | 0.01* | 5.99 | 0.008* | 4.53 | 0.02* |
| cbw | 2.78 | 0.08 | 1.99 | 0.16 | 2.75 | 0.09 | 2.99 | 0.07 |

#### Hippocampus

An ANOVA analysis of hipppocampal volume did not show an effect of genotype, but a significant effect of sex (F(1,23)=19.0, p=0.0002). There were significant effects within all the genotypes APOE2 (t=-2.4, p=0.02), APOE3 (t=-2.9, p=0.008), APOE4 (t=-2.3, p=0.03). Differences between males and females were significant within APOE2 mice (t=-2.4, p=0.02), APOE3 mice (t=-2.9, p=0.008), and APOE4 mice (t=-2.3, p=0.03).

FA only showed a significant effect of genotype (F(2,23)=4.1, p=0.03), and a trend for the genotype by sex interaction (F(2,23)=2.4, p=0.1). Significant differences were found only within groups of females: APOE2 versus APOE3 (t=2.7, p=0.03), and APOE2 versus APOE4 (t=3.2, p=0.009). Differences between groups of males were not significant. Sex differences between animals of the same genotype were borderline significant for APOE4 mice, but not for the other genotypes (t=-1.7, p=0.1).

The clustering coefficient showed a significant effect of genotype (F(2,23)=6.9, p=0.004), and a trend for sex (F(1,23)=2.8, p=0.1). Interestingly, differences were significant between males of APOE2 and APOE3 genotypes (t=-2.7, p=0.03). There were no differences between males and females of the same genotype.

#### Caudate Putamen

The ANOVA analysis for the caudate putamen showed a singificant effect of genotype (F(2,23)=13.8, p=0.0001). Differences between groups of females were significant for APOE2 versus APOE4 mice (t=-2.8, p=0.02), and between APOE3 and APOE4 mice (t=-4.4, p<0.001). Differences between groups of males were only significant between APOE3 and APOE4 mice (t=-2.9, p=0.02). Differences between males and females within the same genotypes were borderline significant for APOE4 mice (t=1.7, p=0.1).

FA analyses indicated a significant effect of genotype (F(2,23)=3.9, p=0.03), and differences within groups of females were borderline significant for APOE2 versus APOE4 (p=2.4, p=0.06). Also differences between male and female AP0E44 mice were borderline significant (t=-1.9, p=0.07).

We did not detect significant differences in the degree and clustering coeffcient of the caudate putamen.

#### Fimbria and Fornix

The fimbria volume was borderline significant for the interaction term genotype by sex (F(2,23)=2.6, p=0.1). The FA only showed a trend for genotype (F(2,23)=2.9, p=0.1). We did not detect significant differences in the clustering coeffcient. The ANOVA analyses for the fornix did not show an effect for genotype and sex.

#### Internal Capsule

We found no differences in the volume of the internal capsule, however the FA showed a significant effect of genotype (F(2,23)=5.5, p=0.01), as well as sex (F(1,23)=17.7, p=0.0003). Within groups of females, differences between APOE2 and APOE4 mice were significant (t=3.1, p=0.01), and showed a trend between APOE2 and APOE3 mice (t=2.5, p=0.05). We found no significant differences between groups of males of different genotypes. An analysis within genotypes showed differences between APOE3 mice of different sexes (t==2.5, p=0.02), and between APOE4 mice of different sexes (t=-3.9, p=0.002).

The degree of connectivity showed significant effects for genotype (F(2,23)=6, p=0.008), sex (F(1,23)=5.7, p=0.03) and the interaction of genotype by sex (F=3.6, p=0.045). Within groups of females we identified differences between APOE2 and APOE3 mice (t=-3.8, p=0.003), and APOE3 and APOE4 mice (t=4.1, p=0.001). These differences were not seen within groups of males. Differences between males and females were only identified for APOE3 mice (t=3.5, p=0.002).

Similar differences as for the degree of connectivity we noticed for the clustering coefficient, showing significant effects for genotype (F(2,23)=4.5, p=0.02), sex (F(1,23)=5.0, p=0.03), but the interaction between genotype and sex was not significant. Within groups of females we identified differences between APOE2 and APOE3 mice (t=-3.4, p=0.007), and between APOE3 and APOE4 mice (t=3.1, p=0.01). These differences did not persist within groups of males. Differences between males and females were only found for APOE3 mice (t=2.9, p=0.008).

#### Cerebellar White Matter

For volume, we only identified a trend for the genotype effect (F(2,23)=2.8, p=0.08), but a significant effect of sex (F(1,23)=18.9, p=0.0002), and for the genotype by sex interaction (F(2,23)=7.9, p=0.002). Post hoc tests identified significant differences between females of APOE2 and APOE3 genotypes (t=-3.5, p=0.005), and between females of APOE3 and APOE4 genotypes (t=3.1, p=0.01). For male mice diferences were significant between APOE2 and APOE4 mice (t=3.1, p=0.01). For mice of the same genotypes sex differences were significant for APOE3 (t=4.9, p=5.6*10^-5^), and also for APOE4 mice (t=3.2, p=0.004).

FA showed a significant interaction between genotype and sex (F(2,23)=5.3, p=0.01). Within groups of females APOE2 and APOE3 showed significant differences (t=3.8, p=0.002), and a trend for APOE3 and APOE4 mice (t=-2.4, p=0.06). Sex differences were identified for APOE3 mice only.

The degree of connectivity only showed a trend for the effect of genotype (F(3,23)=2.7, p=0.1), and this paralelled our results for the clustering coefficient, which showed a trend for the genotype effect (F(2,23)=3.0, p=0.07).

In conclusion, genotype differences were noted for the volume of the caudate putamen, the FA of the hippocampus, caudate putamen, fimbria and fornix, and the connectivity of the hippocampus and internal capsule.

#### Spatial navigation trajectory shape as a function of imaging parameters

We built linear models for the AWN during the two probes for the hippocampus, caudate putamen, and their connecting tracts, as well as the cerebellar white matter and assessed the significance of the relationships between AWN and regional imaging metrics for all mice (**Table 3**).

**Table 3.** Main ANOVA results on the linear models predicting AWN based on MRI metrics.

| Day5 |  |  |  |
| --- | --- | --- | --- |
| Region | Metric | F | p |
| CPu | Volume | 9.86 | 0.006* |
| fi | Volume | 17.81 | 0.0006* |
| Hc | FA | 5.23 | 0.02* |
| CPu | FA | 4.51 | 0.048* |
| ic | FA | 7.77 | 0.02* |
| fx | FA | 4.003 | 0.06 |

| Day8 |  |  |  |
| --- | --- | --- | --- |
| Region | Metric | F | p |
| CPu | Volume | 42.25 | 5.45E-06* |
| fx | Volume | 10.61 | 0.005* |
| Hc | FA | 10.15 | 0.005* |
| fx | FA | 5.39 | 0.03* |
| ic | FA | 12.61 | 0.002* |
| fi | DEG | 2.96 | 0.1 |
| fx | DEG | 5.35 | 0.03* |
| ic | DEG | 7.35 | 0.02* |
| fi | CLUS | 5.78 | 0.03* |
| fx | CLUS | 4.85 | 0.04* |
| cbw | CLUS | 6.06 | 0.03* |
| CPu | FA | 3.8 | 0.07 |

We examined whether the relationships between AWM and imaging metric differed for mice of different genotypes and sexes (**Table 4**, **Figure 7** and **Figure 8**).

**Figure 7.**
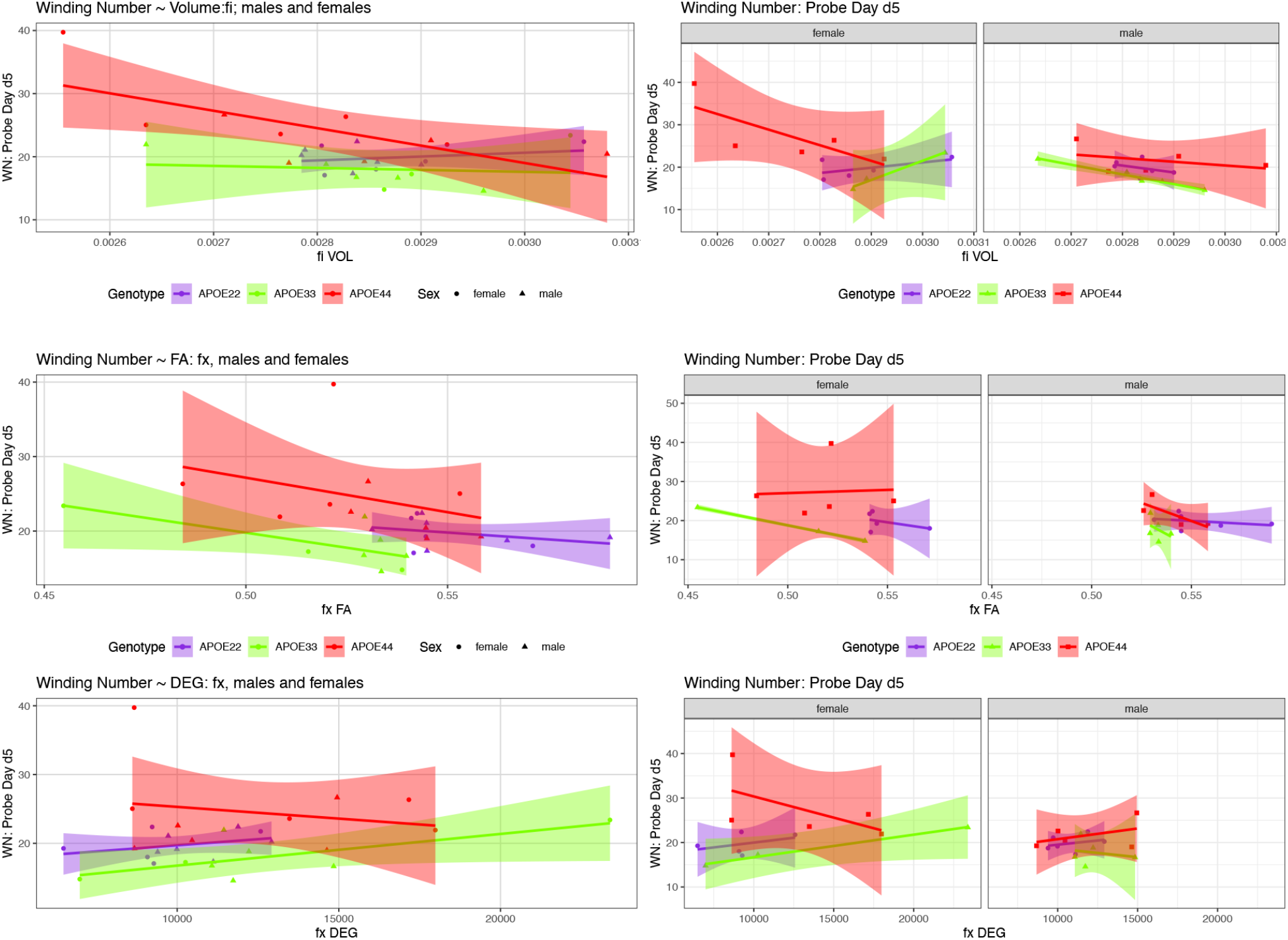
The day 5 AWN~ MRI metrics models. Slopes were different for the fimbria volume between APOE2 and APOE4 females (p=0.03), and APOE3 versus APOE4 females (p=0.008), and there was a trend for APOE3 versus APOE4 slopes (p-0.07). There was also a significant difference in slopes between males and females with APOE3 genotype (p=0.01). There were also significant differences between APOE3 and APOE4 females in the slopes for fornix FA (p=0.01), and degree of connectivity (p=0.03). There was also a trend for the slope differences between APOE3 and APOE4 females for the internal capsule volume (p=0.1, not shown).

**Figure 8.**
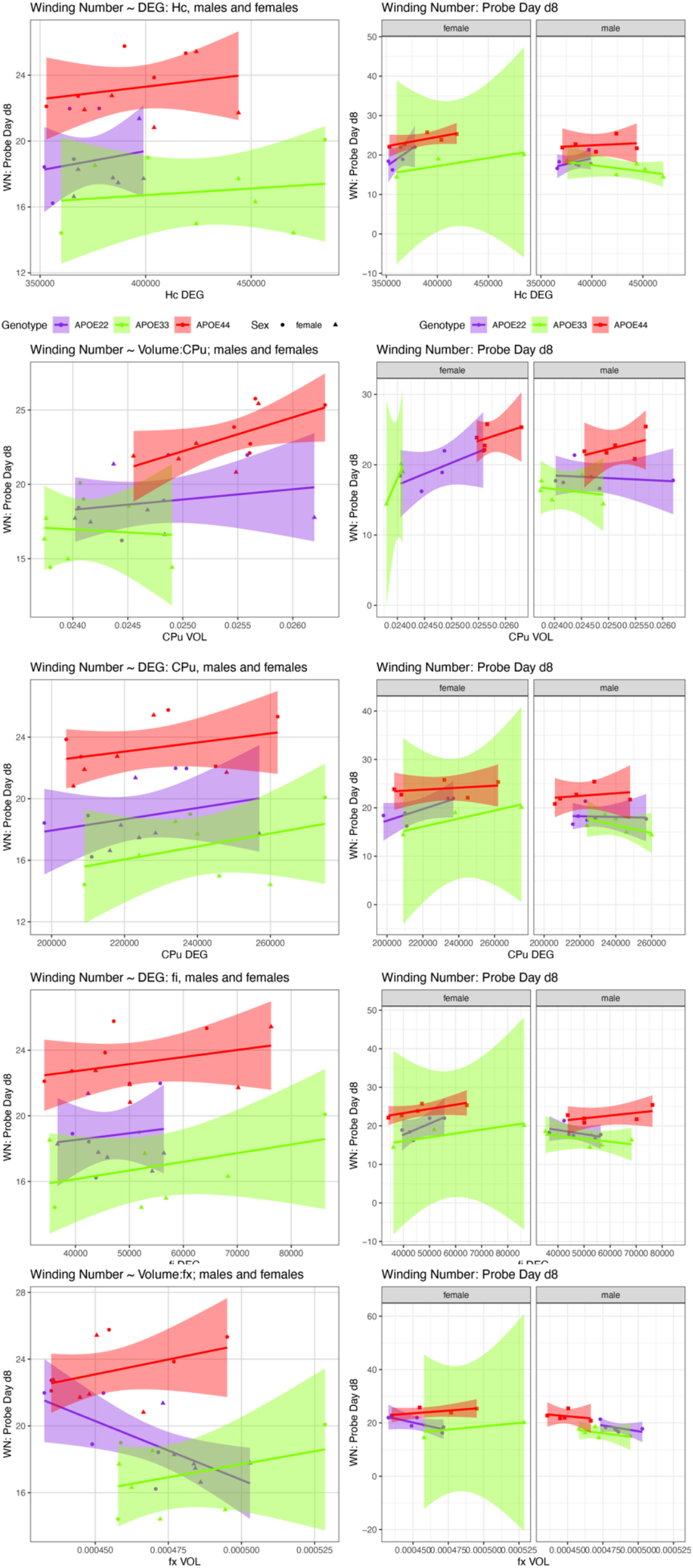
The day 8 AWN~ MRI metrics models. APOE3 mice showed differences between males and females in the slopes for the hippocampus degree of connectivity, caudate putamen volume, and degree of connectivity, while males and female APOE2 mice were different for the fimbria degree of connectivity. The fornix volume showed differences between females APOE2 and APOE3 (p=0.04), and there was a trend for APOE2 versus APOE4 differences (p=0.06).

**Table 4.**
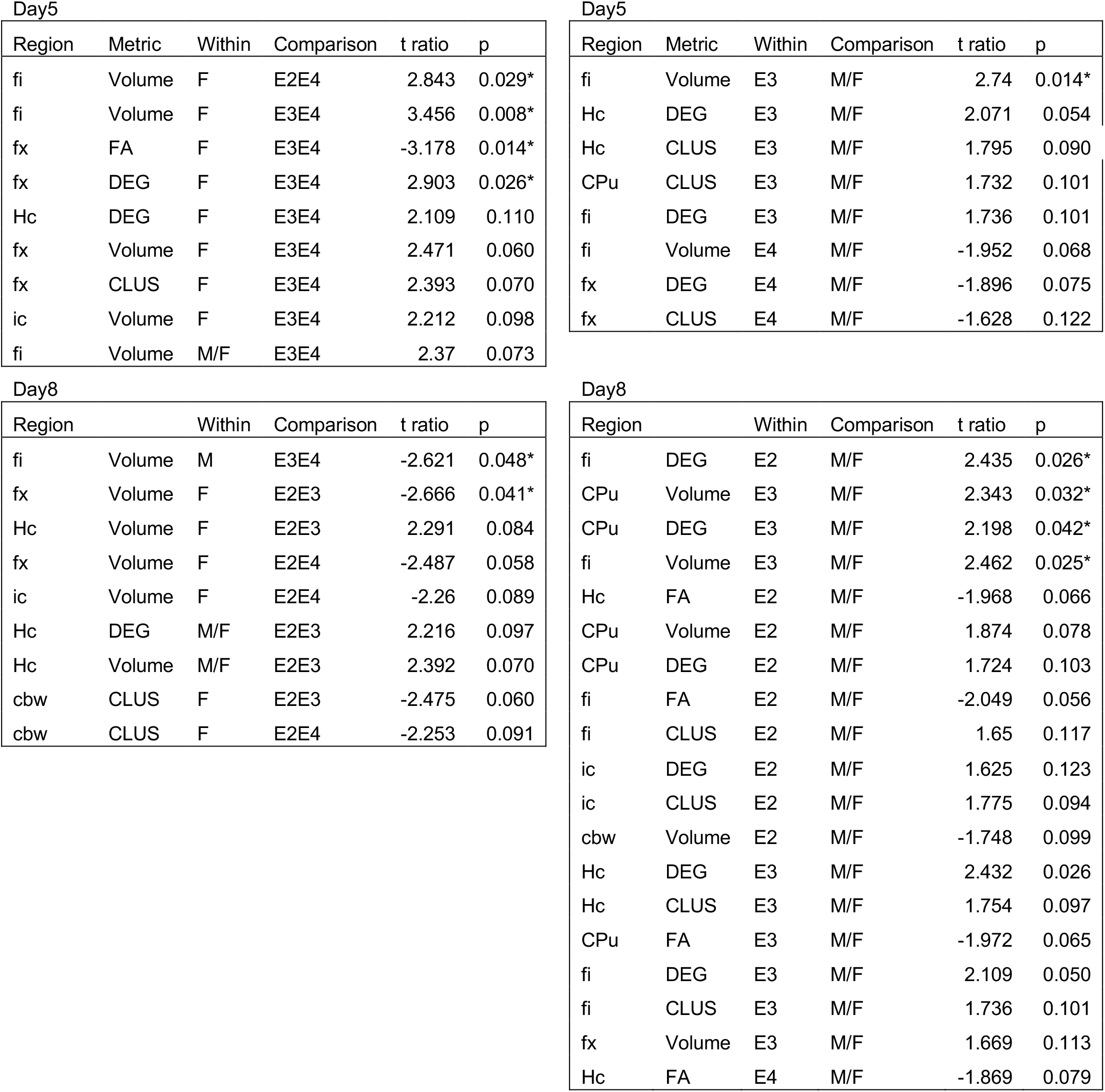
Summary of AWN~MRI metrics comparisons. APOE22: E2, APOE33: E3, APOE44: E4; F: female; M: male.

| Region | Metric | Within | Comparison | t ratio | p |
| --- | --- | --- | --- | --- | --- |
| fi | Volume | F | E2E4 | 2.843 | 0.029* |
| fi | Volume | F | E3E4 | 3.456 | 0.008* |
| fx | FA | F | E3E4 | -3.178 | 0.014* |
| fx | DEG | F | E3E4 | 2.903 | 0.026* |
| Hc | DEG | F | E3E4 | 2.109 | 0.110 |
| fx | Volume | F | E3E4 | 2.471 | 0.060 |
| fx | CLUS | F | E3E4 | 2.393 | 0.070 |
| ic | Volume | F | E3E4 | 2.212 | 0.098 |
| fi | Volume | M/F | E3E4 | 2.37 | 0.073 |

Day5
| Region | Metric | Within | Comparison | t ratio | p |
| --- | --- | --- | --- | --- | --- |
| fi | Volume | E3 | M/F | 2.74 | 0.014* |
| Hc | DEG | E3 | M/F | 2.071 | 0.054 |
| Hc | CLUS | E3 | M/F | 1.795 | 0.090 |
| CPu | CLUS | E3 | M/F | 1.732 | 0.101 |
| fi | DEG | E3 | M/F | 1.736 | 0.101 |
| fi | Volume | E4 | M/F | -1.952 | 0.068 |
| fx | DEG | E4 | M/F | -1.896 | 0.075 |
| fx | CLUS | E4 | M/F | -1.628 | 0.122 |

Day8
| Region |  | Within | Comparison | t ratio | p |
| --- | --- | --- | --- | --- | --- |
| fi | Volume | M | E3E4 | -2.621 | 0.048* |
| fx | Volume | F | E2E3 | -2.666 | 0.041* |
| Hc | Volume | F | E2E3 | 2.291 | 0.084 |
| fx | Volume | F | E2E4 | -2.487 | 0.058 |
| ic | Volume | F | E2E4 | -2.26 | 0.089 |
| Hc | DEG | M/F | E2E3 | 2.216 | 0.097 |
| Hc | Volume | M/F | E2E3 | 2.392 | 0.070 |
| cbw | CLUS | F | E2E3 | -2.475 | 0.060 |
| cbw | CLUS | F | E2E4 | -2.253 | 0.091 |

| Region |  | Within | Comparison | t ratio | p |
| --- | --- | --- | --- | --- | --- |
| fi | DEG | E2 | M/F | 2.435 | 0.026* |
| CPu | Volume | E3 | M/F | 2.343 | 0.032* |
| CPu | DEG | E3 | M/F | 2.198 | 0.042* |
| fi | Volume | E3 | M/F | 2.462 | 0.025* |
| Hc | FA | E2 | M/F | -1.968 | 0.066 |
| CPu | Volume | E2 | M/F | 1.874 | 0.078 |
| CPu | DEG | E2 | M/F | 1.724 | 0.103 |
| fi | FA | E2 | M/F | -2.049 | 0.056 |
| fi | CLUS | E2 | M/F | 1.65 | 0.117 |
| ic | DEG | E2 | M/F | 1.625 | 0.123 |
| ic | CLUS | E2 | M/F | 1.775 | 0.094 |
| cbw | Volume | E2 | M/F | -1.748 | 0.099 |
| Hc | DEG | E3 | M/F | 2.432 | 0.026 |
| Hc | CLUS | E3 | M/F | 1.754 | 0.097 |
| CPu | FA | E3 | M/F | -1.972 | 0.065 |
| fi | DEG | E3 | M/F | 2.109 | 0.050 |
| fi | CLUS | E3 | M/F | 1.736 | 0.101 |
| fx | Volume | E3 | M/F | 1.669 | 0.113 |
| Hc | FA | E4 | M/F | -1.869 | 0.079 |

##### Hippocampus

###### Day 5

There was a significant effect of the FA on the absolute winding number for day 5 (F(1,7)=7.3, p=0.02). Differences between female groups were borderline significant (F(1,7)=4.8, p=0.07).

There was a trend for the interaction between the degree of connectivity and genotype (F(2,17)=2.5, p=0.1). Also, there was a trend within males (F(1,10)=4.6, p=0.06). There was a trend for the slopes differences between APOE3 and APOE4 females (t=1.1, p=0.1, and between males and females APOE3 carriers (t ration =2.1, p=0.05). There was a significant effect for the clustering coefficient as a predictor for the winding number at day 5 for males (F(1,10)=5.1, p<0.05), a trend for APOE3 females (t ratio 1.58, p=0.1), and for the differences between slopes for males and female APOE3 (t=1.8, p=0.09).

###### Day 8

There was a significant effect of FA (F(2,17)=10.2, p=0.005), and a trend for sex (F(1,17)=3.07, p=0.1), and the interaction FA by sex (F(1,17)=4.22, p0.06). The efect was significant within females (F(1,7)=12.1, p=0.01). There was a trend for slope differences between females and males of APOE2 genotypes (t ratio=-2, p=0.07), and APOE4 (t ratio=-2, p=0.08). There was only a trend for the degree of connectivity interaction by sex F(1,17)=2.9, p=0.1. The slopes were different between males and females APOE3 mice (t ratio 2.4, p=0.03).

##### Caudate Putamen

###### Day 5

There was a significant effect of volume on the winding number at day 5 (F(1,17)=9.9, p=0.006). The effect within females was characterized by F(1,7)=5.5, p=0.05.

There was also a significant effect of FA on the absolute winding number (F(1,17)=4.5, p<0.05), and for the interaction of FA with sex (F(1,17)=4.47, p<0.05). There was a significant effect of FA on AWN within females (F(1,7)=6.5, p=0.04), and a trend for slopes differences between males and females with APOE4 genotypes (t ratio =-2.0, p=0.06).

The degree of connectivity showed a significant effect as a predictor within males (F(1,10)=7, p=0.02). This was paralelled by the clustering coeffcient showing as a significant predictor within males (F(1,10)=10.4, p=0.009). There was a trend for slopes differences between females and males of APOE3 genotypes (t ratio =1.7, p=0.1).

###### Day 8

There was a significant effect of the CPu volume on the AWS F(1,17)=42.3, p=5*10^-6^), and the effect was significant both within males (F(1,10)=8.2, p=0.02), and within females F(1,7)=34.7, p=0.006). There was a difference between slopes for APOE3 males and females (t ratio=2.3, p=0.03).

There was a significant effect of FA within females only (F(1,7)=11.8, p=0.01).

There was a significant effect of the degree of connectivity both withing males (F(1,10)=7.7, p=0.02), and females (F(1,7)=6.5, p=0.04). The slopes were different within APOE3 mice (t ratio=2.2, p=0.04), and showed a trend within APOE2 mice (t ratio=1.7, p=0.1).

There was a significant difference in slopes for the AWN versus CPu clustering coefficient between APOE3 male and female mice (t ratio=2.4, p=0.03).

##### Fimbria

###### Day 5

There was a significant effect of the fimbria volume on the AWN (F(1,17)=17.8, p=0.0006), as well as a significant interaction between the volume, genotype, and sex (F(2,17)=6.0, p=0.01). The effect was significant in females (F(1,7)=10.7, p=0.01), as well as for the interaction for fimbria volume by genotype (F(2,7)=5.0, p<0.05). There was a trend for slopes diffrences for APOE3 and APOE4 mice (t ratio=2.4, p=0.07). These differences were significant between groups of females for APOE2 versus APOE4 mice (t ratios=2.8, p=0.03), and for APOE3 and APOE4 female mice (t ratio=3.5, p=0.008). Differences were significant between female and male APOE3 mice (t ration =2.7, p=0.01), and showed a trend between female and male APOE4 mice (t=-2, p=0.07). The slopes were different than 0 for APOE4 mice (t=-3.2, p=0.006), and in particular for APOE4 females (t ratio=-3.7, p=0.002), and showed a trend for APOE3 females (t=2.1, p=0.05), and males (t=-1.9, p=0.08).

The FA interaction with genotype was significant (F(2,17)=5.1, p=0.04). There was a trend for slopes for APOE3 (t=-1.6, p=0.1) and APOE4 mice (t =1.6, p=0.1), and for slope differnces between APOE3 and APOE4 females (t =-2.2, p=0.1).

For the degree of connectivity we only found a trend between females and males with APOE3 genotypes (t=1.7, p=0.1). The clustering coefficient interaction by sex also showed a trend (t=3.2, p=0.09).

###### Day 8

There was a significant difference between the slopes for volume and AWS between APOE3 and APOE4 mice (t ratio=-2.6, p=0.04).

For the degree of connectivity there was a significant interaction with sex (F(1,17)=6.3, p=0.02), and there were significant differences between males and females of APOE2 (t ratio=2.4, p=0.03); and a trend for APOE4 (tratio=2.1, p=0.05).

There was a significant effect of the clustering coefficient (F(1,17)=5.8, p=0.03).

##### Fornix

###### Day 5

There was a trend for the volume as a predictor of AWN for females APOE3 versus APOE4, t =2.5, p=0.06), and for female versus male APOE4 carriers (t=-1.7, p=0.1), and this was similar with the degree of connectivity for APOE4 females versus males (t=-1.9, p=0.08).

###### Day 8

There was a significant effect for the fornix volume (F(1,17)=10.6, p=0.005), and a trend for the interaction of fx volume by genotype F(2,17)=2.5, p=0.1. This was significant in males (F(1,10)=22.0, p=0.001). The slopes were different between APOE2 and APOE3 female mice (t ratio=-2.7, p=0.04) and there was a trend between APOE2 and APOE4 mice (t ratio=-2.5, p=0.06).

There was a significant effect for fx FA (F(1,17)=5.4, p=0.04, with a trend for APOE3 females (p=0.1).

There was a significant effect for the degree of connectivity F(2,17)=24.3, p=1.1*10^-5^), and this was significant within females (F(1,7)=6.3, p=0.04). The clustering coefficient was also significant (F(1,17)=4.9, p=0.04). This was significant within females (F(1,7)=6.5, p=0.04), with a trend for APOE3 females (p=0.1).

##### Internal Capsule

###### Day5

There was significant interaction of the volume by genotype (F(2,17)=3.7, p=0.04) and a trend for the slope diffrences between APOE3 and APOE4 females (t=2.2, p=0.1).

There was a significant effect of FA (F(1,17)=7.8, p=0.01). There was a trend for the clustering coeffcient slope differences between APOE4 females and males (t=1.7, p=0.1).

###### Day8

There was a significant effect of the FA (F(1,17)=12.6, p=0.002), with a trend within females (F(1,7)=5.2, p=0.06). There was a significant effect of the degree of connectivity within males (F(1,10)=7.3, p=0.02).

##### Cerebellar White Matter

###### Day 5

We found no effects of the cerebellum white matter on the winding number at day 5.

###### Day8

There was an effect of the clustering coefficient F(1,17)=6.1, p=0.02), with a trend for differnces between APOE2 and APOE3 (t =-2.3, p=0.1), and APOE2 and APOE4 (t =-2.5, p=0.06)

Our comparison of the models’ slopes revealed differences between groups of females with different APOE carriage, both at day 5 (**Figure 7**), and day 8 (**Figure 8**), emphasizing the role of the fornix and fimbria, and suggesting that these major players may interact with other brain regions forming more complex network that determine spatial navigation. Sex differences were also noted, including in the control genotype APOE3 in these circuits, suggesting possible sex modulation of genetic risk for AD.

## 4 Discussion

The major known genetic risk for sporadic, or late onset AD is linked to the APOE gene, and it is conferred by the presence of APOE4 allele. Studying human subjects, or animal models with APOE4 carriage is thus an important strategy for discovering early biomarkers predictive of abnormal aging. However, in cognitively normal subjects, APOE4 is not always associated with an increased risk of cognitive deterioration, suggesting that APOE4 effects on structural and/or clinical progression only become evident in mild cognitive impairment (MCI) and AD (Haller et al. 2017). Still, several studies have shown spatial navigation/orientation deficits in AD, and some indicated that these changes are present in MCI patients and even in cognitively healthy APOE4 carriers (Coughlan et al. 2018). It is important to answer the question whether APOE4 carriers at risk for AD perform spatial navigation tasks differently from APOE2 and APOE3 carriers. If true, spatial navigation and orientation might provide novel cognitive evaluation metrics for prodromal or incipient AD, as sensitive and specific markers of the disease (Coughlan et al. 2018). Rodents provide tools to model AD at prodromal stages, and test novel interventions to remove pathologies, and slow cognitive decline; thus we were motivated to explore spatial learning, memory and navigation strategies in mouse models with different genetic risk for AD.

Our premise lies in knowing that humans and also rodents use preferentially one of two navigational strategies. A spatial strategy (Packard, Hirsh, and White 1989; Iaria et al. 2003) relies on forming relationships between landmarks in the environment and orienting oneself in relation to those landmarks. This process requires the ability to form cognitive maps of the environment and the flexibility to derive a direct path to a target during navigation. The spatial strategy is subserved by the hippocampus (Morris et al. 1982). In contrast, a response strategy involves learning a series of stimulus-response associations, e.g. the pattern of left and right turns from a given starting position. This strategy relies on the caudate putamen, and is inflexible, in that it does not require generating a de novo, direct path to a target location (Packard, Hirsh, and White 1989) during navigation.

The most popular method to assess spatial learning and memory in rodents is the MWM, and several adaptations of this test have been proposed and adopted in human research. The memory and learning processes are usually characterized by distance and time measures to a hidden platform, or the distance and time spent in the target quadrant during learning trials, or probe tests. Few publications have characterized the swim patterns, and this was usually done by assigning the swim path, according to its shape (Brody and Holtzman 2006), (Brabazon et al. 2017), (Zhao et al. 2012), into a small number of discrete categories: direct, chaining, scanning, etc. (Janus 2004). The proportions of time, or distance spent in each of these categories was then compared.

In this work we have introduced a new metric, the AWN, to the battery of tests and metrics used for assessing the cognition of mouse models of neurological conditions, such as AD. This provides a quantitative way to describe the continuous curve that is the swim trajectory, during goal directed spatial navigation. Our analyses showed that this metric is robust to noise, and can be used to compare and better separate relatively small groups of mice, based on their spatial navigation strategies. The AWN was sensitive to genotype and sex, discriminating APOE3 mice as having simpler trajectories during the learning process relative to APOE2 and APOE4 mice, and this effect was strongest in females. The probe trials revealed that APOE4 mice had more complex, loopier trajectories during memory tests.

We have examined whether differences in memory, and spatial navigation strategies were accompanied by imaging and connectivity changes, and how these metrics were related to the AWN. This is because proper memory function requires structural and functional connections of networks (Linden 2007; Piccoli et al. 2015), e.g. involving the dorsal hippocampus (Hc) for spatial memory, and the ventral hippocampus (Hc) for emotional memory (Fanselow and Dong 2010b; Fanselow and Dong 2010a). In rodents, the dorsal Hc and subiculum form a critical network with the anterior cingulate, that mediates processes such as learning, memory, and navigation.

Our results showed that mouse models representing different levels of genetic risk for Alzheimer’s disease performed differently in the spatial memory tests, as assessed with the Morris Water Maze. We added to the existing body of knowledge the observation that swim paths differ with genotypes not just in length but also in shape. We introduced a new metric through the absolute winding number, which gives insight into spatial navigation strategy differences, is robust to noise, and showed differences between females. Moreover, the absolute winding number discriminated APOE3 carriers during learning trials, as they have simpler trajectories relative to APOE2 and APOE4 carriers, which are more similar, and these differences are due to females. During probe trials administered at 1 hour after the end of learning, the absolute winding number discriminated APOE4 mice relative to APOE2 and APOE3 carriers, as these two groups had more similarly shaped trajectories. Our data on the spatial search strategy tested three days after the end of the learning trials suggest a genotype “dose” dependent effect, and this was particularly apparent in females, while APOE4 males were differentiated relative to APOE2 and APOE3 males, that had more similar search strategies.

These behavioral changes were accompanied by differences in the volume of the caudate putamen, but not the hippocampus. We did however find significant changes in the hippocampal FA, its degree of connectivity, and clustering coefficient. These underline the roles of hippocampal microstructural properties and connectivity, and suggest such changes may precede overt neurodegeneration, i.e. atrophy. The hippocampal degree of connectivity and clustering coefficient did discriminate between APOE2 and APOE3 mice. The degree of connectivity was also different between APOE2 and APOE4 mice.

Besides the hippocampus, microstructural changes were found in the caudate putamen and fornix, and there was a trend for the internal capsule. Changes in the degree of connectivity were found for the hippocampus and internal capsule, with a trend for the cerebellar white matter. The clustering coefficient was different for the hippocampus, and showed a trend for the fornix, internal capsule, and cerebellar white matter. The clustering coefficient for the hippocampus differentiated APOE2 versus APOE3 mice, and there was a trend for the fornix, internal capsule, and cerebellar white matter. These results suggest that carriage of different APOE alleles results in different connectivity for regions involved in circuits related to spatial navigation, learning and memory, as well as the associated motor task execution. Together, our results support the importance of the fornix in rodent spatial navigation, in agreement with evidence from human studies (Hodgetts et al. 2020).

As we hypothesized that imaging metrics could help predict changes in the spatial trajectory shape, the AWN on day 5 found that hippocampal FA, as well as the caudate putamen volume and FA, differed significantly among the three APOE genotypes. There was an effect of internal capsule FA. We also found that the slopes of AWN~ fimbria volume were significantly different for APOE3 versus APOE4 females, and for APOE2 versus APOE4 females. Our results denoted that different strategies were used by APOE4 females. Interestingly the AWS~ internal capsule only had a zero slope for APOE4 mice (data not shown), suggesting these mice may rely more on striatal circuits to accomplish their goal oriented navigation task.

At 3 days after the last learning trial (day 8), we found stronger relationships between the imaging metrics and the AWN. The hippocampal FA, caudate putamen volume, as well as the fornix volume were also significant. Importantly, these data support the role of the fornix in determining the shape of the swim path, or spatial navigation strategy, as all metrics were significant (volume, FA, degree of connectivity, and clustering coefficient). The internal capsule FA and degree of connectivity were also significant. Regions for which connectivity was a predictor of the AWN at 3 days after the last learning trial were the fimbria, fornix, and cerebellum white matter. In summary our data support the role of the fornix in spatial memory and navigation, and demonstrates involvement of other regions, including the caudate putamen, and cerebellar white matter.

Due to our limited sample sizes, and the fact that we only investigated a small set of regions, we were unable to dissect whole circuits, or the different roles of these structures in different genotypes. However, our data showed slope differences for the AWN~fimbria model within females: APOE2 vs APOE4 (p= 0.03); and for APOE3 vs APOE4 females (p= 0.0080). At day 8 there were slope differences between males of APOE3 and APOE4 genotypes (t=-2.6, p<0.05). This suggests that different circuits, or different contributions of the same circuits in spatial navigation in mice with different APOE genotypes, and of different sexes. Further studies should investigate the association between vulnerable brain circuits and cognitive traits, in particular to reveal sex differences.

Ours is not a comprehensive study to dissect the role of vulnerable circuits in spatial navigation, learning and memory. Rather it is proposing a hypothesis, based on a subselection of brain regions and connections, in particular those involving the hippocampal (allocentric) and striatal (egocentric, and procedural) based circuits. These circuits are likely to interact in spatial navigation, and our data suggest that the presence of different APOE alleles plays role (Goodroe, Starnes, and Brown 2018). This is important in the context of AD related changes in spatial memory, as it may point to specific pathways (Neuner et al. 2017). The use of this metric in a full brain analysis will likely provide important new leads in our quest to understand the early changes of APOE-related vulnerability and mechanisms, and to reveal early biomarkers.

## Author Contributions

AB, DL, CAC, and DD conceived the study, devised methods, wrote and edited the manuscript. NB generated the mouse lines, suggested ideas and analyses, edited the manuscript; and together with CAC and CLW participated in the interpretation of the behavioral and imaging data. AN, and AB performed experiments. AB, AN, RJA, JAS, DL performed analyses.

## Acknowledgments

This work was supported by RF1 AG057895, and R01 AG066184, U24 CA220245, RF1 AG070149, R01 MH118927, and the Bass Connections program at Duke. We are grateful to the members of the Duke Radiology and Duke Neurology Departments for generously sharing advice, support and resources needed for the experiments, in particular Gary Cofer, Lucy Upchurch, Brian Mace, Viviana Cantillana, and Joan G Wilson. We are grateful to NIH and the Bass Connections program for supporting our research.

## Data Availability Statement

The datasets analyzed for this study can obtained from the first authors. Code is shared at https://github.com/AD-Decode/awn.

## Contribution to the Field

We studied animals modeling genetic risk for late onset Alzheimer’s disease using behavior and MR imaging. We introduce a new metric, the absolute winding number, to characterize the spatial search strategy, through the shape of the swim path. The absolute winding number better differentiated APOE3 carriers, through their straighter swim paths relative to APOE2 and APOE4 genotypes. This novel metric was sensitive to sex differences, supporting increased vulnerability in APOE4 females. We hypothesized differences in spatial memory and navigation strategies are linked to differences in brain networks, and showed that different genotypes have different reliance on the hippocampal and caudate putamen circuits, pointing to a role for white matter connections. Our results support a departure from a hippocampal centric to a brain network approach, and open new avenues for identifying regions linked to increased risk for AD, before overt disease manifestation. Further exploration of novel biomarkers based on spatial navigation strategies may enlarge the windows of opportunity for interventions. The proposed framework will be significant in dissecting vulnerable circuits associated with cognitive changes in prodromal Alzheimer’s disease.

